# Structural basis of lipid-mediated gating in a two-pore domain potassium channel

**DOI:** 10.1101/2025.11.12.688043

**Authors:** Fumiya K. Sano, Go Kasuya, Kohei Yamaguchi, Takehiro Suzuki, Daichi Yamanouchi, Kana M. Hashimoto, Masataka Inoue, Kazuhiro Sawada, Ryuichiro Ishitani, Kenjiro Yoshimura, Hisato Hirano, Yuzuru Itoh, Koichi Nakajo, Naoshi Dohmae, Yoshiaki Kise, Osamu Nureki

## Abstract

TOK1 is the first identified member of the K2P channel family and contains additional N- and C-terminal domains, displaying a configuration distinct from that of canonical K2P channels. Although recent advances in structural studies of K2P channels have elucidated their architectures, the structural basis of TOK1 has remained unknown, limiting our understanding of its unique configuration. Here, we present the cryo-electron microscopy (cryo-EM) structure of TOK1, unveiling its distinctive domain architecture. Furthermore, the structures of TOK1 in three distinct states provide mechanistic insights into its regulation through lipid binding and dissociation. Phosphorylation of TOK1 induces the formation of an additional lipid-binding site, leading to channel inactivation. Conversely, upon activation, the phospholipid dissociates, allowing ion permeation. Our comprehensive study, integrating cryo-EM structural analysis, molecular dynamics simulations, electrophysiological recordings, and mass spectrometry, elucidates the distinctive features of TOK1, an atypical K2P channel, and provides a framework for understanding lipid-mediated regulation within this family.

## Introduction

Two-pore domain potassium (K2P) channels constitute a unique family of potassium channels defined by distinctive structural and functional features^1–4^. Each K2P channel contains two pore-forming domains, and two subunits assemble into a dimeric architecture to form a functional potassium channel. K2P channels are responsible for background or “leak” K^+^ conductance, thereby contributing to the maintenance of the resting membrane potential and the regulation of cellular excitability^1–5^. Their activities are regulated by a wide range of physicochemical stimuli — including mechanical stretch, pH, temperature, and lipid composition — and each member has evolved to specialize in sensing distinct stimuli^6–8^. Recent structural studies have revealed the structural basis of K2P channels. Specifically, the structures of several members such as TWIK, TRAAK, and TASK have provided insights into their overall architectures and common gating mechanisms^9–20^. Notably, the structural bases of lipid-mediated gating mechanisms have been reported for TREK1 and THIK1^1^^2,17–20^.

TOK1, an outwardly rectifying potassium channel, represents the first identified member of the K2P channel family and is widely conserved among fungi^21^. The topology predicted from the amino acid sequence of TOK1 is markedly different from that of mammalian K2P channels. TOK1 contains an additional four transmembrane helices at the N-terminus and an intracellular soluble domain at the C-terminus (**Supplementary Fig. 1a**). Furthermore, TOK1 lacks the conserved extracellular cap domain that is commonly observed in mammalian K2P channels (**Supplementary Fig. 1a–c**). Despite a long history of study using diverse approaches^22,23^, the detailed functional mechanism of TOK1 has not yet been revealed, thereby underscoring the need for structural analysis.

In this study, we present the cryo-electron microscopy (cryo-EM) structure of the TOK1 channel, revealing its unique overall architecture. Furthermore, the structures of TOK1 in three distinct states elucidate how phospholipids regulate channel activity involving dynamic conformational changes. By integrating molecular dynamics simulations, mass spectrometry, and electrophysiological experiments, this study substantially deepens our understanding of TOK1, a unique member of the K2P channel family.

## Results

### Overall architecture of TOK1

To investigate the unique configuration predicted from the amino acid sequence, we performed cryo-EM structural analysis of TOK1 (**Supplementary Fig. 1a,b**). First, we purified *Saccharomyces cerevisiae* TOK1 expressed in HEK293S GnTI^−^ cells and reconstituted it into a lipid nanodisc (**Supplementary Fig. 2a,b**). Then, we performed cryo-EM analysis of TOK1 and determined its structure at a resolution of 2.6 Å (**Fig. 1a–f, Supplementary Fig. 2c–h, Supplementary Table 1**). The resulting density map showed a local resolution of 2.0–3.5 Å, allowing confident model building for most regions except for the N- and C-termini and several loop regions (**Supplementary Fig. 2f**).

**Fig. 1.**
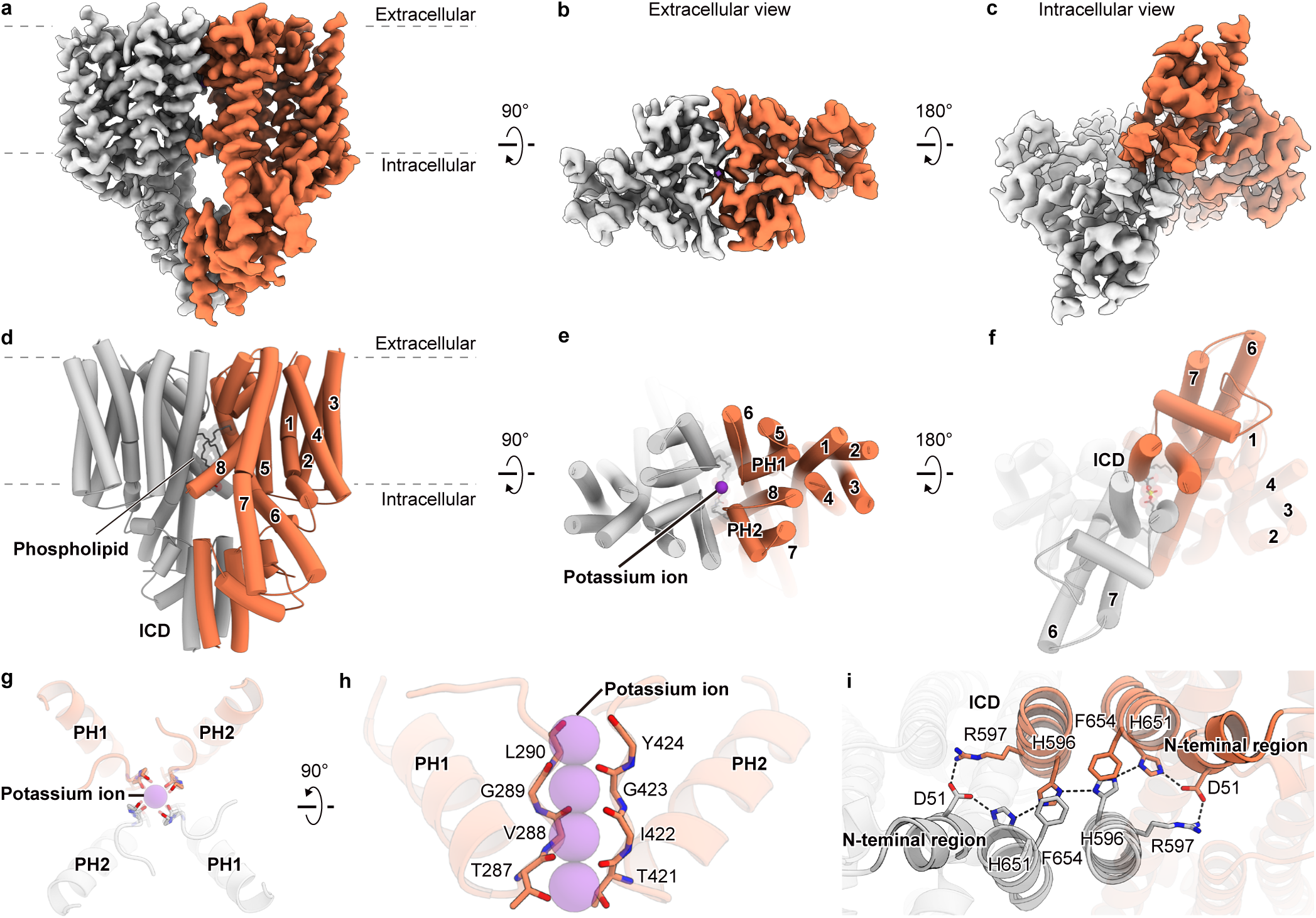
Overall architecture of TOK1. **a–c.** Cryo-EM density map of TOK1 viewed from the side (**a**), extracellular (**b**), and intracellular (**c**) orientations. **d–f.** Cylinder models of TOK1 viewed from the side (**d**), extracellular (**e**), and intracellular (**f**) orientations. **g, h.** Selectivity filter of TOK1 viewed from the extracellular (**g**) and side (**h**) orientations. The main chains of key residues are shown in stick representation (**g, h**). **i.** Intracellular domain (ICD) of TOK1 viewed from the intracellular orientation. Key interacting residues at the protomer interface are shown in stick representation (**i**). The two protomers are shown in orange and light gray, respectively (**a–i**).

Similar to many other two-pore channels, TOK1 forms a dimer that creates a central ion-conducting pore (**Fig. 1d–f**)^9–20^. However, as predicted from its amino acid sequence, TOK1 exhibits an architecture distinct from that of other two-pore channels (**Supplementary Fig. 1a,b**). Here, we describe the TOK1 structure by dividing it into three regions—the N-terminal helices, pore-forming helices, and intracellular domain—and briefly outline the features of each (**Supplementary Fig. 1a**). First, the four N-terminal transmembrane helices are positioned distally from the ion-conducting pore and form a stable helical bundle that likely contributes to the overall stability of TOK1 (**Fig. 1d–f, Supplementary Fig. 1a**). Second, TMs 5–8 exhibit an architecture similar to that of canonical two-pore domain potassium channels^9–20^. Around the pore helices (PH1 and PH2), two selectivity filter motifs orient their main-chain oxygen atoms toward the center of the ion-conducting pore (**Fig. 1e,g,h**). However, unlike other two-pore channels, TOK1 lacks a cap structure between TM5 and TM6, and no domain swapping—in which the first transmembrane helix is exchanged between protomers—was observed (**Fig. 1d,e, Supplementary Fig. 1b,c**). Third, the C-terminal intracellular domain (ICD) is positioned between the two protomers, flanked by the extended TM5 and TM6 helices (**Fig. 1f**). At the center of the ICD, two helices from each subunit assemble into a four-helix bundle. Together with the N-terminal region, this bundle forms an extensive network of polar interactions and π-stacking that strongly stabilizes dimer formation (**Fig. 1i**). Overall, TOK1 adopts a unique configuration distinct from that of canonical two-pore domain potassium channels.

### A phospholipid blocks ion permeation

Interestingly, a phospholipid molecule is bound at the ion-conducting pore, obstructing ion permeation and defining this structure as being in the closed state (**Fig. 2a**). To ensure that this phospholipid is not an artifact of the nanodisc environment, we also determined the structure of TOK1 solubilized in glyco-diosgenin (GDN) (**Supplementary Fig. 3, Supplementary Table 1**). A similar phospholipid density was observed at the same position, indicating that phospholipid binding is an intrinsic feature of TOK1 (**Supplementary Fig. 1d,e**). The phospholipid is recognized through distinct interactions involving its head group and acyl chains, respectively. First, the head group of the phospholipid is located on the twofold (C2) symmetry axis and forms salt bridges with R323 (**Fig. 2a**). Notably, considering that TOK1 is a potassium channel, the presence of a positively charged residue, R323, near the ion-conducting pore appears rather unusual, suggesting that phospholipid binding serves as an intrinsic regulatory mechanism. The distal portion of the head group shows poorly defined density. We modeled this phospholipid as POPC; however, it is likely that multiple phospholipid species can bind at this site (**Fig. 2c, Supplementary Fig. 2d**). Second, the two acyl chains of the phospholipid are positioned between the protomers and are tightly packed by numerous hydrophobic residues on TM6 and TM8 (**Fig. 2b**). Thus, the phospholipid binds above the ion-conducting pore and negatively regulates TOK1 function by inhibiting ion permeation.

**Fig. 2.**
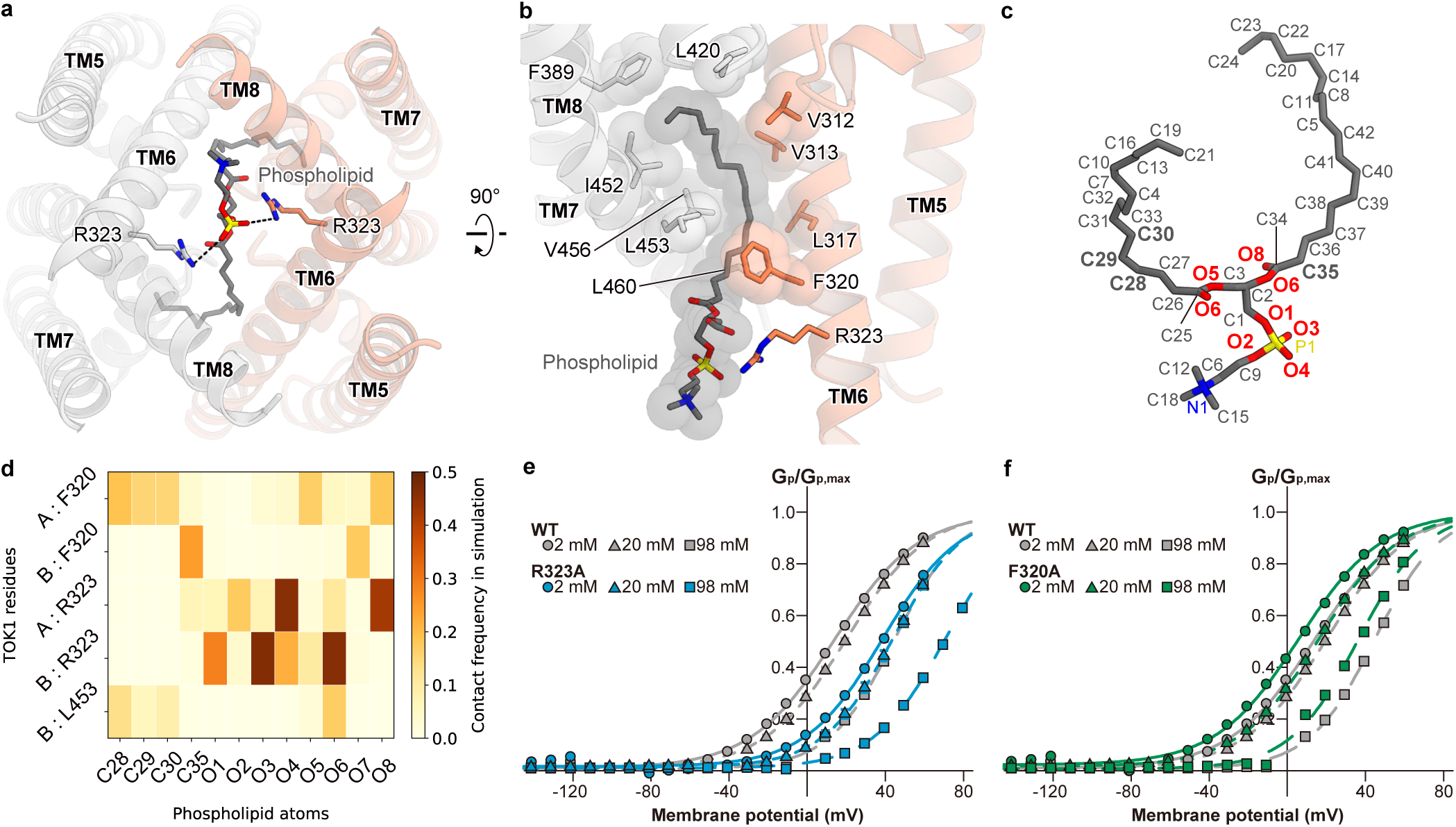
Binding mode of the phospholipid. **a, b.** Binding mode of the phospholipid viewed from the intracellular (**a**) and side (**b**) orientations. Key residues for phospholipid recognition are shown in stick and CPK representations. The two protomers are shown in orange and light gray, respectively (**a, b**). **c.** Stick representation of the phospholipid with atom name labels. Labels of key interacting residues are shown in bold (**c**). **d.** Contact frequencies between TOK1 and the phospholipid in molecular dynamics simulations. The horizontal axis represents the atoms of the phospholipid, and the vertical axis represents the residues of TOK1. The prefixes A and B indicate each protomer, respectively. Residues (TOK1) or atoms (phospholipid) with a contact frequency below 15% are omitted. The heat map represents the combined data from three independent replicate simulations (**d**). **e, f.** Peak conductance–voltage (G_p_–V) relationships of wild-type (WT) and R323A mutant TOK1, and of WT and F320A mutant TOK1, under external K^+^ concentrations of 2 mM (filled circles), 20 mM (filled triangles), and 98 mM (filled squares). The G_p_–V relationships were fitted with a single Boltzmann function. Symbols and bars represent mean ± s.e.m. (n = 8)(**e, f**).

To further investigate the phospholipid recognition by TOK1, we performed molecular dynamics (MD) simulations. Using the cryo-EM structure of TOK1 as the initial model, 250-ns simulations were conducted, and the overall structure remained stable throughout three independent production runs (**Supplementary Fig. 4a–c**). We analyzed the contact frequencies between TOK1 and the phospholipid in the MD simulations, which revealed several key residues for phospholipid recognition. First, the head group and nearby oxygen atoms frequently contacted R323, suggesting critical recognition through polar interactions, as observed in the cryo-EM structure (**Fig. 2c,d**). In addition, several carbon atoms of the acyl chains showed frequent contacts with F320 (**Fig. 2c,d**). F320 appears to play a dominant role in the recognition of the acyl chains, which are highly flexible and thus recognized through multiple weak hydrophobic interactions. These observations are consistent with the cryo-EM findings and collectively provide valuable insight into the mechanism underlying lipid recognition by TOK1.

To examine the effect of phospholipids on the gating of TOK1, we conducted electrophysiological experiments using *Xenopus* oocytes. Specifically, two-electrode voltage clamp (TEVC) recordings were performed to assess the activity of wild-type TOK1 (TOK1^WT^) and several of its mutants. TOK1^WT^ exhibited pronounced outward rectification and a clear dependence on extracellular K^+^ concentration, consistent with previous reports^21,24,25^, confirming that our experimental system functioned properly (**Supplementary Fig. 5a,f**). Unexpectedly, the R323A mutant, which is involved in recognizing the phospholipid head group, exhibited a depolarizing shift of approximately 20 mV in the G–V curve, suggesting reduced channel activity (**Fig. 2e**). In contrast, the F320A mutant, which is involved in recognizing the phospholipid acyl chains, exhibited a hyperpolarizing shift in the G–V curve, suggesting enhanced channel activity due to destabilization of the bound phospholipid (**Fig. 2f**). In any case, both mutations clearly altered the channel properties of TOK1, suggesting that phospholipids regulate its gating.

### Inhibition mechanism through phosphorylation

TOK1 is negatively regulated by Hog1, a mitogen-activated protein (MAP) kinase activated under high-osmolarity conditions^26^. As a first step toward understanding how phosphorylation controls channel gating, we identified the phosphorylation sites of TOK1. Constitutively active Pbs2 (a MAP kinase kinase for Hog1), wild-type Hog1, and TOK1 were individually purified and then mixed in the presence of ATP, followed by incubation at 37 °C for 1 h. The reaction mixture was separated by gel filtration, and the fractions corresponding to TOK1 were treated with peptidase and subsequently analyzed by mass spectrometry (**Supplementary Fig. 6**). As a result, the phosphorylation sites of TOK1 were found to be mainly clustered in the loop region between TM8 and the ICD, specifically at S474, S548, S550, S551, S556, S557, S568, S569, S577, and T641 (**Fig. 3a**).

**Fig. 3.**
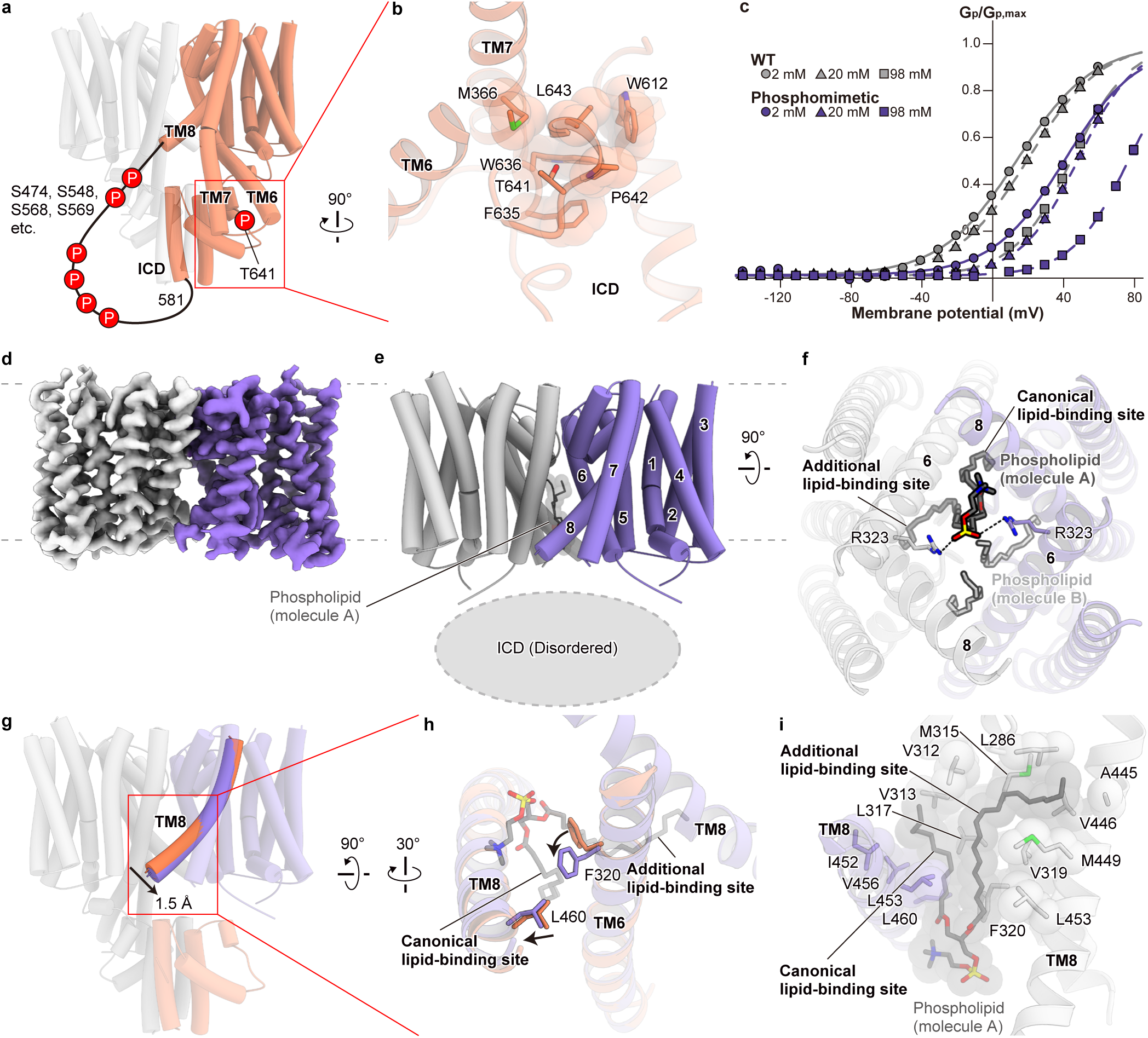
Mechanism of inhibition through phosphorylation. **a.** Representation of the identified phosphorylation sites in TOK1. **b.** Phosphorylation site T641 in the intracellular domain (ICD) of TOK1. **c.** Peak conductance–voltage (G_p_–V) relationships of WT and Phosphomimetic mutant TOK1, under external K^+^ concentrations of 2 mM (filled circles), 20 mM (filled triangles), and 98 mM (filled squares). The G_p_–V relationships were fitted with a single Boltzmann function. Symbols and bars represent mean ± s.e.m. (n = 8)(**c**). **d, e.** Cryo-EM density map and cylinder model of TOK1^phosphomimetic^ viewed from the side orientations. **f.** Binding mode of the phospholipid viewed from the intracellular orientation. The two bound phospholipid molecules are shown in dark gray and light gray, respectively. For clarity, the headgroup of one molecule is omitted (**f**). **g, h.** Conformational changes around TM6 and TM8 induced by the phosphomimetic mutation, viewed from the side (**g**) and intracellular (**h**) orientations. The flip of F320 forms an additional lipid-binding site (**h**). **i.** Phospholipid recognition mechanism of TOK1^phosphomimetic^.

To investigate the mechanism by which phosphorylation negatively regulates channel activity, we conducted structural studies. First, we generated a phosphomimetic mutant of TOK1 (TOK1^phosphomimetic^) in which the identified serine and threonine phosphorylation sites were substituted with glutamate residues. As expected, TOK1^phosphomimetic^ exhibited a depolarizing shift in the G–V curve (**Fig. 3c**), confirming that the phosphomimetic mutation negatively regulates TOK1 in a manner similar to phosphorylation. We then determined the cryo-EM structure of TOK1^phosphomimetic^ at 3.0 Å resolution (**Fig. 3d,e, Supplementary Fig. 7, Supplementary Table 1**). Surprisingly, the ICD of TOK1^phosphomimetic^ was completely disordered. Considering that most of the residues subjected to the phosphomimetic mutations are located in the loop region between TM8 and the ICD, the disorder observed in the ICD of TOK1^phosphomimetic^ is likely attributable to T641, the only site within the ICD (**Fig. 3a,b**). In TOK1^WT^, the side chain of T641 is oriented toward a cage composed of hydrophobic bulky residues from TM7 and the ICD (**Fig. 3b**). This cage is not large enough to accommodate a phosphate group, suggesting that phosphorylation at T641 likely induces substantial structural rearrangements of ICD. Overall, TOK1 is suggested to undergo pronounced structural rearrangements upon phosphorylation, resulting in disorder of the ICD.

Notably, TOK1^phosphomimetic^ exhibits an alternative lipid-binding mode distinct from that of the wild type. As in the wild type, an acyl chain was observed at the canonical lipid-binding site, located on the TM6–TM8 interface between the opposing protomers (**Fig. 3f, Supplementary Fig. 1d,f**). Importantly, another acyl chain was observed at an additional lipid-binding site located on the intra-protomer TM6–TM8 interface. This distinct phospholipid-binding mode arises from structural rearrangements in TM6 and TM8. As a result of ICD disorder, TM8 of TOK1^phosphomimetic^ adopts a relatively vertical orientation with respect to the plasma membrane (**Fig. 3g**). This structural rearrangement of TM8 induces a flip of the side chain of F320 on TM6 (**Fig. 3h**). The resulting space is occupied by the acyl chain of a phospholipid, defining an additional lipid-binding site. At the additional lipid-binding site, the acyl chain is stabilized through hydrophobic interactions with multiple residues (**Fig. 3i**). The presence of this extra lipid-binding site potentially facilitates phospholipid binding from an entropic perspective, thereby stabilizing the closed state of the channel. Alternatively, the binding of the acyl chain at the additional lipid-binding site may simply be stronger than that at the canonical lipid-binding site. Taken together, these results suggest that phosphorylation of TOK1 induces ICD disorder and triggers structural rearrangements in TM6 and TM8, leading to the formation of an additional lipid-binding site and thereby negatively regulating channel activity.

### Activation mechanism of TOK1

Several mutations of TOK1 that markedly inhibit yeast growth have been reported^24,25^. Considering that knockout of TOK1 has little effect on yeast growth, these mutations are suggested to promote channel opening and cause severe cellular damage. We tested the electrophysiological properties of the TOK1^T322I^ mutant, one of these mutants. As expected, we observed a hyperpolarizing shift in the G–V curve, indicating enhanced channel activity (**Fig. 4a**). To investigate the activation mechanism of TOK1, we determined the cryo-EM structure of the TOK1^T322I^ mutant at a resolution of 2.9 Å (**Fig. 4b,c, Supplementary Fig. 8, Supplementary Table 1**). Surprisingly, no phospholipids were observed in the ion-conducting pore of TOK1^T322I^ (**Supplementary Fig. 1g**). Furthermore, TM5–TM8 undergo conformational changes from the closed state to ensure a pore radius sufficient for potassium ion permeation, thereby defining this structure as the open state (**Fig. 4d,e**).

**Fig. 4.**
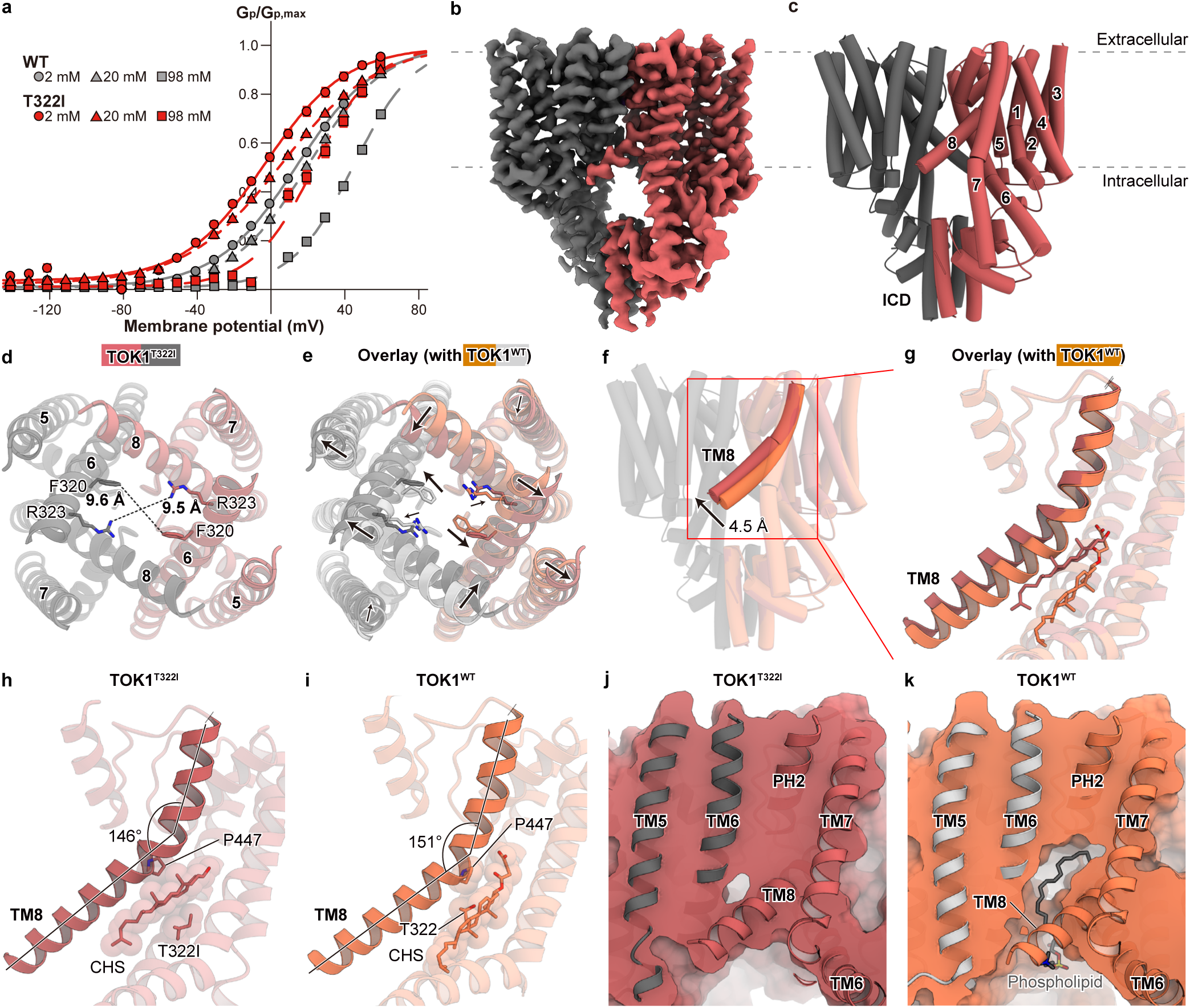
Activation mechanism of TOK1. **a.** Peak conductance–voltage (G_p_–V) relationships of WT and T322I mutant TOK1, under external K^+^ concentrations of 2 mM (filled circles), 20 mM (filled triangles), and 98 mM (filled squares). The G_p_–V relationships were fitted with a single Boltzmann function. Symbols and bars represent mean ± s.e.m. (n = 8)(**a**). **b, c.** Cryo-EM density map and cylinder model of TOK1^T322I^ viewed from the side orientations. **d, e.** Ion-conducting pore of TOK1^T322I^ (**d**) and its overlay with TOK1^WT^ (**e**), viewed from the intracellular orientation. **f–i.** Conformational changes induced by the T322I mutation. TOK1^T322I^ and TOK1^WT^ exhibit distinct binding modes of cholesterol or its analogue, cholesterol hemisuccinate (CHS)(**g–i**). The angle of TM8 was measured using the Cα atoms of T431, A442, and S464 (**h, i**). **j, k.** Molecular surface representation of TOK1^T322I^ and TOK1^WT^. The phospholipid-binding site observed in TOK1^WT^ is not observed in TOK1^T322I^ (**j, k**).

The open-state structure of the TOK1^T322I^ clarifies the activation mechanism of this channel. In TOK1^T322I^, TM8 adopts a relatively horizontal orientation with respect to the plasma membrane, which arises from the distinct binding mode of cholesterol or its analogue, cholesterol hemisuccinate (CHS) (**Fig. 4f,g**). T322I pushes TM8 toward the extracellular side with CHS positioned between, resulting in a pronounced kink in TM8 originating at P447 (**Fig. 4h**). In contrast, in TOK1^WT^, CHS binds at a different position, and the push of TM8 by T322 is limited (**Fig. 4i**). As a result, the phospholipid binding sites observed in TOK1^WT^ are largely occluded in TOK1^T322I^, preventing lipid binding (**Fig. 4j,k**). Thus, the T322I mutation alters the conformation of TM8 via a distinct cholesterol binding mode, resulting in the dissociation of the phospholipid and ultimately leading to gate opening.

We note that two previously identified growth-inhibitory mutations, S330F and V456I, are also strongly suggested to promote channel opening through a similar mechanism, given their locations (**Supplementary Fig. 9a**). This suggests that the activation mechanism revealed in this study is an intrinsic property of TOK1, rather than a phenomenon unique to the T322I mutant.

## Discussion

We determined the cryo-EM structures of the two-pore domain potassium channel TOK1 in the open and closed states. Our structures reveal that a phospholipid bound within the ion-conducting pore negatively regulates channel activity (**Fig. 5**). Moreover, phosphorylation of the intracellular domain (ICD) further suppresses channel opening by forming an additional lipid-binding site (**Fig. 5**). On the other hand, upon activation, TM8 adopts a horizontal orientation relative to the plasma membrane, resulting in the dissociation of the phospholipid and ultimately allowing ion permeation (**Fig. 5**). Integrating cryo-EM structural analysis with molecular dynamics simulations, electrophysiological experiments, and mass spectrometry, this study elucidates the unique activation mechanism of TOK1 mediated by phospholipid binding and dissociation.

**Fig. 5.**
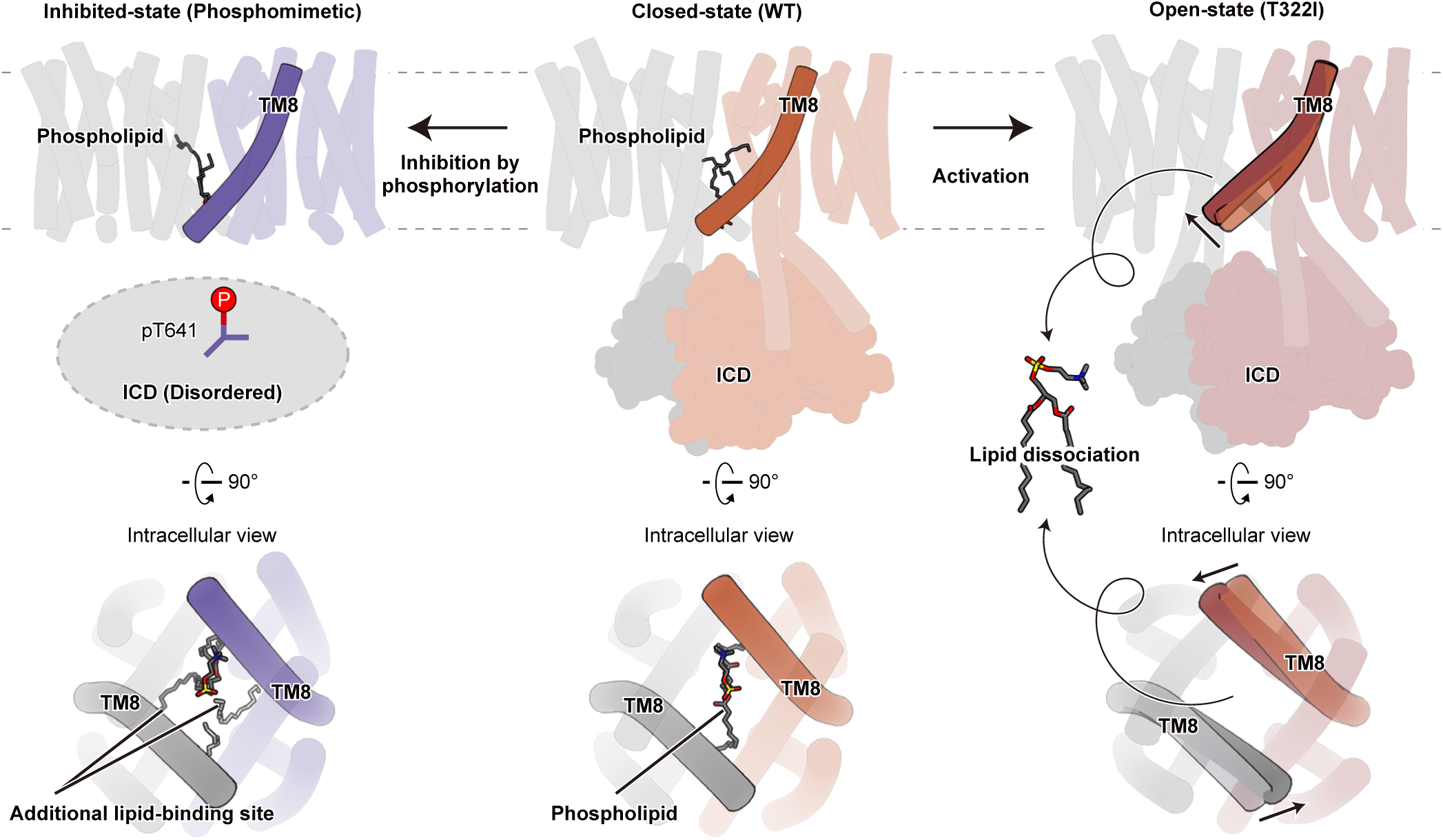
Schematic diagram of the inhibition and activation mechanisms of TOK1. Under resting conditions, a phospholipid binds on the ion-conducting pore, blocking ion permeation (center panel). Phosphorylation of the intracellular domain (ICD) induces its disorder, leading to conformational changes in the transmembrane domain that create an additional lipid-binding site, ultimately further inhibiting ion permeation (left panel). Upon activation, the conformational change of TM8 triggers lipid dissociation, permitting ion permeation (right panel).

As noted above, several growth-inhibitory mutants of TOK1, probably representing open-state mutants, have been identified^25^. Notably, deletions of the ICD have been reported to rescue these growth-inhibitory mutants^24^. These studies strongly suggest that the absence or disorder of the ICD shifts the channel toward the closed state, which is highly consistent with our structural findings.

The ICD also plays a distinct role other than gating regulation. Notably, contact frequency analysis between opposing protomers during MD simulations revealed that not only the pore domain but also the ICD exhibited strong inter-protomer interactions (**Supplementary Fig. 4d,e**). Considering that TOK1 lacks both the Cap domain and the domain swapping of TM5 (TM1 in canonical K2P channels), which are conserved among K2P channels (**Supplementary Fig. 1a–c**), the ICD may also contribute to the dimer stabilization. Thus, the ICD of TOK1 may have two distinct biological roles.

Recent structural studies have demonstrated that several K2P channels are regulated by lipids. For instance, THIK1 is negatively regulated by a fatty acid bound between TM4 and PH1, which stabilizes the channel in an inactive state (**Supplementary Fig. 9b**)^17–20^. Furthermore, TREK1 possesses two phospholipid-binding sites^12^. A lipid bound near the selectivity filter negatively regulates the channel by blocking ion permeation (**Supplementary Fig. 9c**). In contrast, a lipid bound between TM1 and TM4 positively regulates the channel by fixing TM4 in a horizontal conformation relative to the plasma membrane (**Supplementary Fig. 9d**). In both cases, the lipid-binding modes are completely different from those observed in TOK1, highlighting the diverse modes of lipid-mediated regulation within this channel family.

We believe that this study significantly deepens our understanding of not only TOK1 but also two-pore domain potassium channels in general. However, recent studies have revealed that this family, which was previously considered to be merely leak channels, possesses multiple regulatory mechanisms^1–3,6^. These aspects need to be examined in further structural studies.

## Acknowledgements

We thank K. Ogomori, C. Harada, at the University of Tokyo for their technical assistance in structural analysis. This work was supported by grants from the JSPS KAKENHI grant numbers JP23KJ0491 (F.K.S.), JP23H02666 (G.K.), JP21K06786 (K.N.), JP25KJ0866 (K.S.), JP24H01491 (K. Yoshimura), JP25K22482 (K. Yoshimura); the Japan Science and Technology Agency (JST) grant numbers JPMJCR20E2 (O.N.); the Japan Agency for Medical Research and Development (AMED) grant numbers JP25gm7110004 (F.K.S.), JP223fa627001 (F.K.S.), and JP223fa627001 (O.N.); the Platform Project for Supporting Drug Discovery and Life Science Research (Basis for Supporting Innovative Drug Discovery and Life Science Research (BINDS)) from AMED under Grant Number JP23ama121002 (support number 3272) and JP23ama121012.

## Author contributions

F.K.S. prepared the samples for cryo-EM structural analysis with the assistance of K. Yamaguchi, D.Y., K.M.H. and K.S. F.K.S. performed cryo-EM data acquisition, processing, and model building with the assistance of H.H. and Y.I. F.K.S. performed molecular dynamics simulations with the assistance of R.I. G.K. and M.I. performed electrophysiological experiments with the assistance of K.N. and K. Yoshimura. T.S. performed mass-spectrometry analysis with the assistance of N.D. F.K.S., G.K., and K. Yamaguchi wrote the manuscript with input from all authors. G.K., Y.K., and O.N. supervised the research.

## Figures

**Supplementary Fig. 1.**
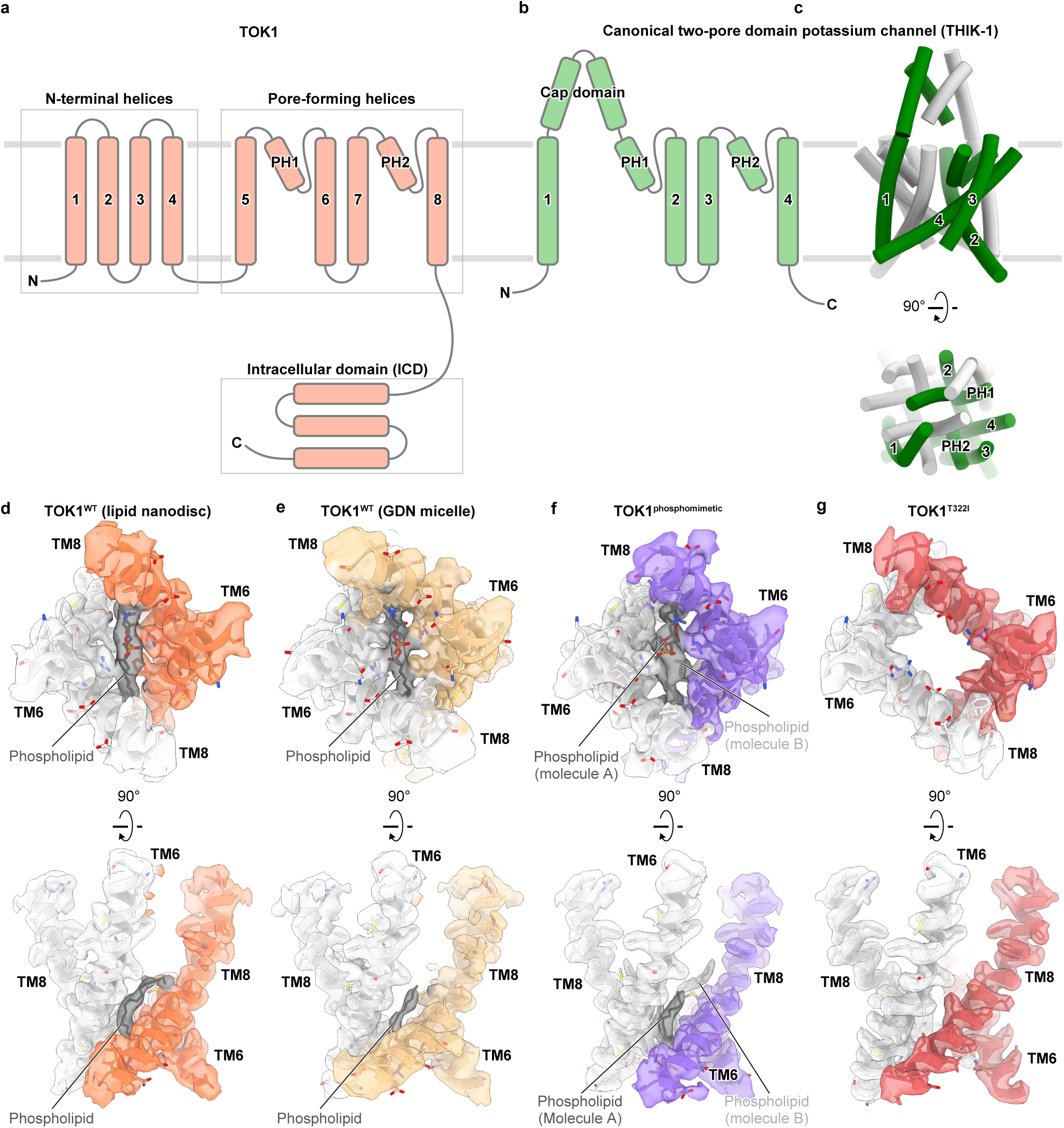
Structural features of TOK1 and canonical two-pore domain potassium channels. **a, b.** Snake models of TOK1 (**a**) and the canonical two-pore domain potassium channel THIK-1 (**b**). **c.** Cylinder model of THIK-1 (PDB 9FT7) viewed from the side (upper panel) and intracellular (lower panel) orientations. The two protomers are shown in orange and light gray, respectively. Domain swapping of TM1 is observed between the opposing protomers (**c**). **d–g.** Map–model overlays of the phospholipid-binding sites above the ion-conducting pore in TOK1^WT^ (lipid nanodisc)(**d**), TOK1^WT^ (GDN micelle)(**e**), TOK1^phosphomimetic^(**f**), and TOK1^T322I^(**g**). Upper and lower panels show intracellular and side views, respectively.

**Supplementary Fig. 2.**
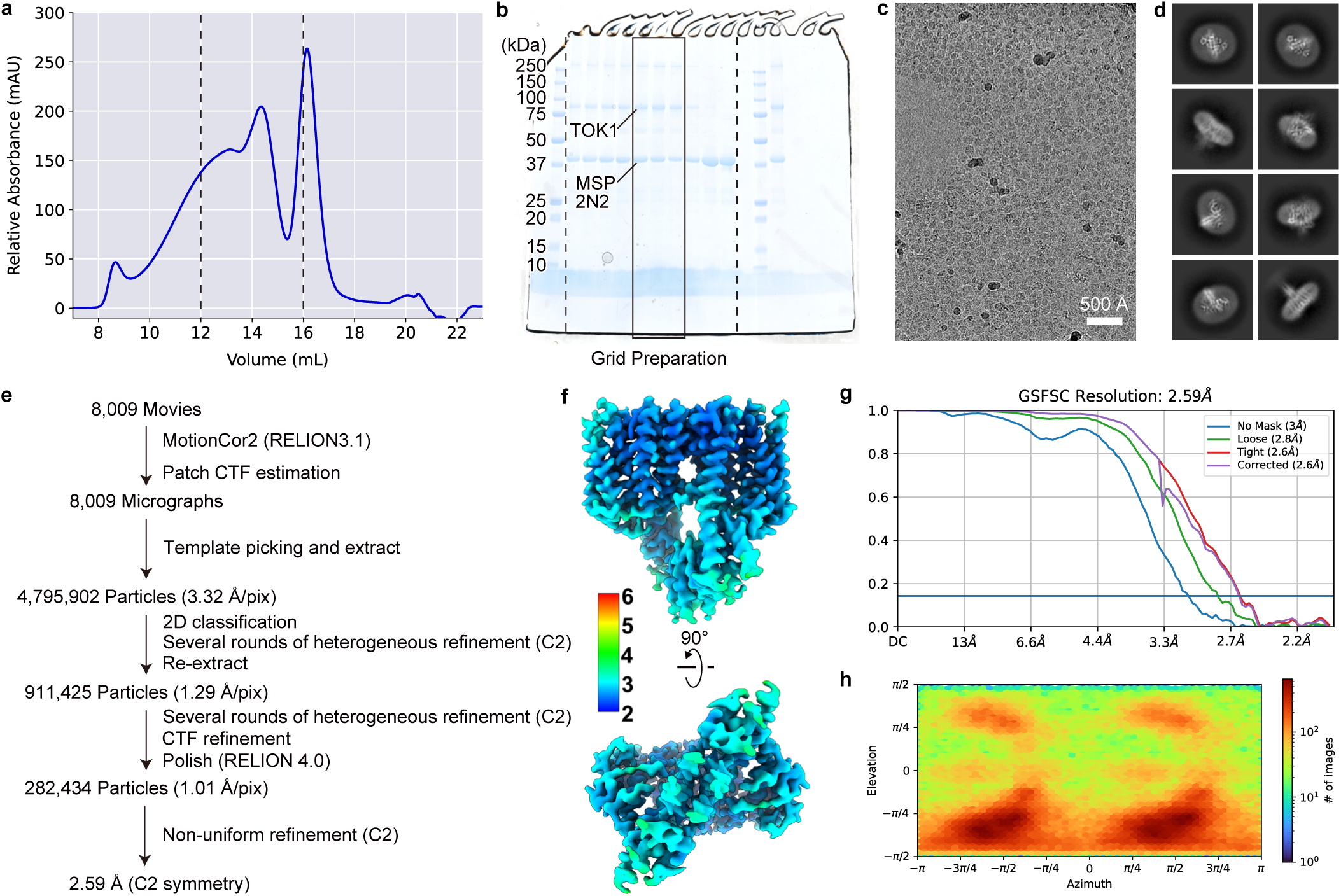
Cryo-EM analysis and map quality of TOK1^WT^ in lipid nanodisc. **a, b.** Size-exclusion chromatography and SDS-PAGE profiles of TOK1^WT^ in lipid nanodisc. **c, d.** Representative cryo-EM micrograph (**c**) and 2D averages (**d**) of TOK1^WT^in lipid nanodisc. **e–h.** Cryo-EM workflow (**e**), local-resolution cryo-EM density map (**f**), Fourier shell correlation (FSC) curve (**g**), and angular distribution plots (**h**) of TOK1^WT^ in lipid nanodisc.

**Supplementary Fig. 3.**
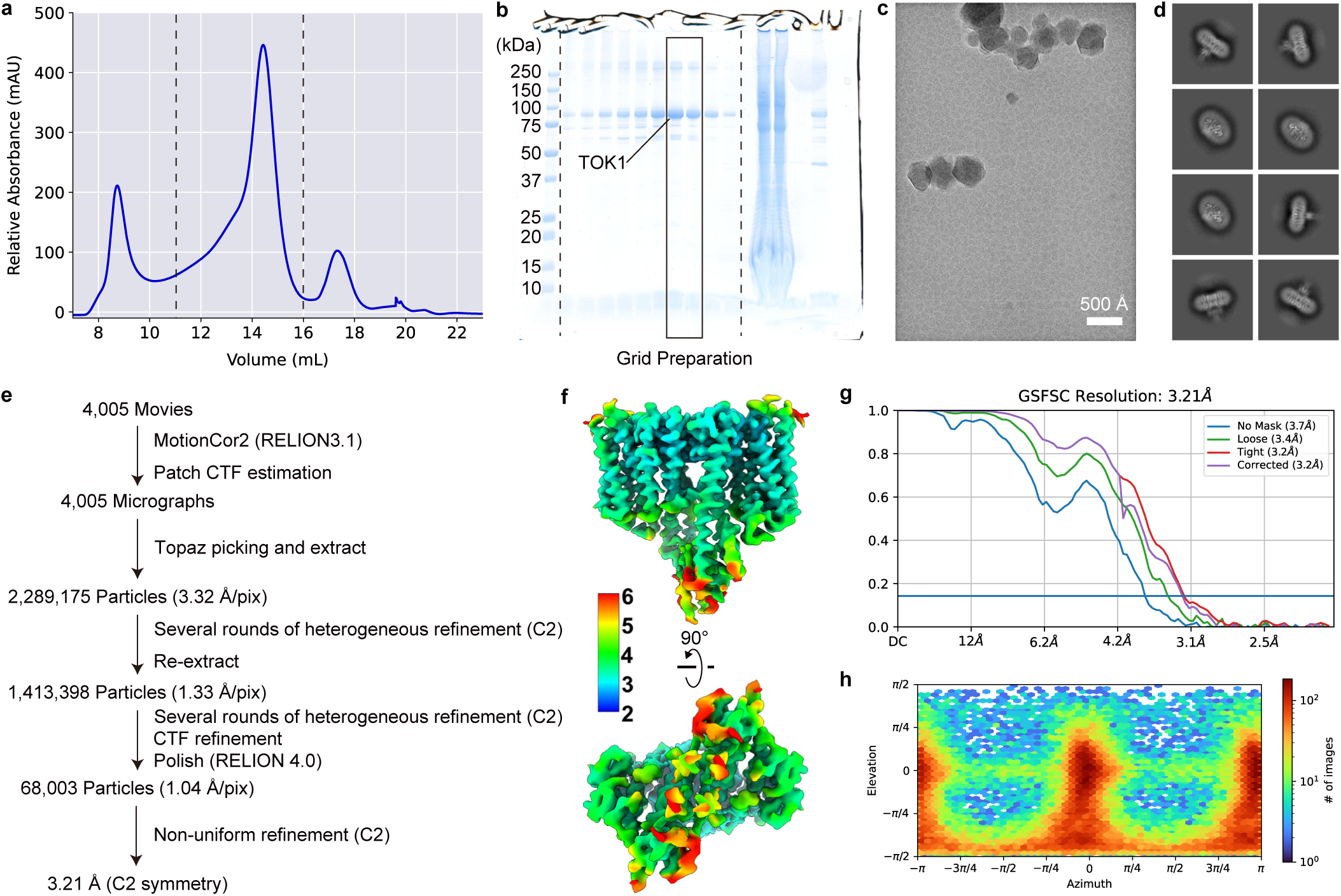
Cryo-EM analysis and map quality of TOK1^WT^ solubilized in GDN. **a, b.** Size-exclusion chromatography and SDS-PAGE profiles of TOK1^WT^ solubilized in GDN. **c, d.** Representative cryo-EM micrograph (**c**) and 2D averages (**d**) of TOK1^WT^ solubilized in GDN. **e–h.** Cryo-EM workflow (**e**), local-resolution cryo-EM density map (**f**), Fourier shell correlation (FSC) curve (**g**), and angular distribution plots (**h**) of TOK1^WT^ solubilized in GDN.

**Supplementary Fig. 4.**
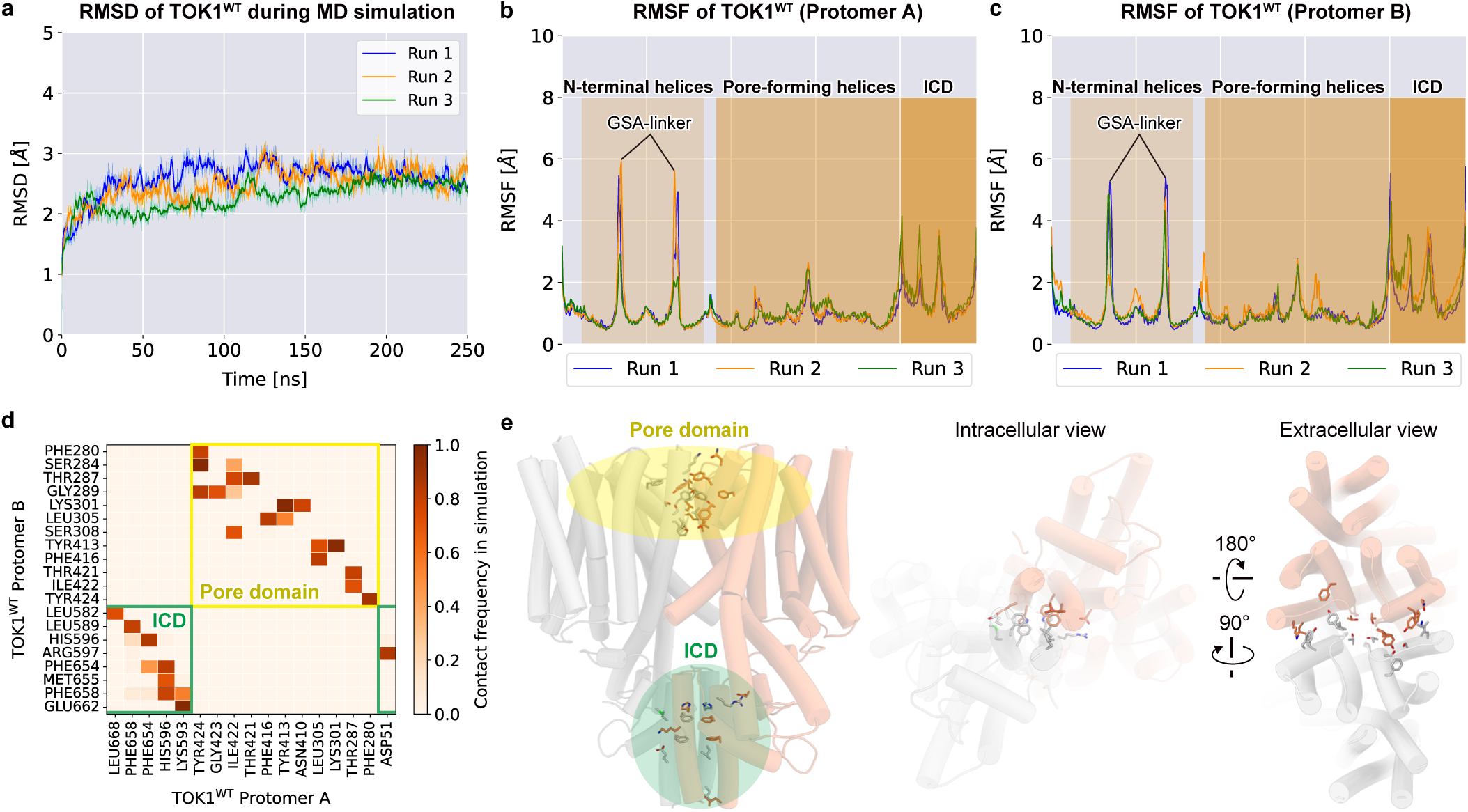
Molecular dynamics simulation of TOK1. **a.** Root-mean-square deviation (RMSD) of Cα atoms of TOK1 during molecular dynamics simulations. Average values, calculated every 10 frames, are shown in bold. **b, c.** Root-mean-square fluctuation (RMSF) of Cα atoms of protomer A (**b**) and protomer B (**c**) of TOK1 during molecular dynamics simulations. The N-terminal helices, pore-forming helices, and intracellular domain (ICD) are highlighted, respectively. **d.** Contact frequencies of inter-protomer interactions in TOK1. The horizontal and vertical axes indicate residues of protomers A and B, respectively. Residues with contact frequencies below 70% are omitted. The heat map shows the combined data from three independent MD simulation replicates (**d**). **e.** Residues that showed high contact frequencies in the MD simulations are displayed as stick models.

**Supplementary Fig. 5.**
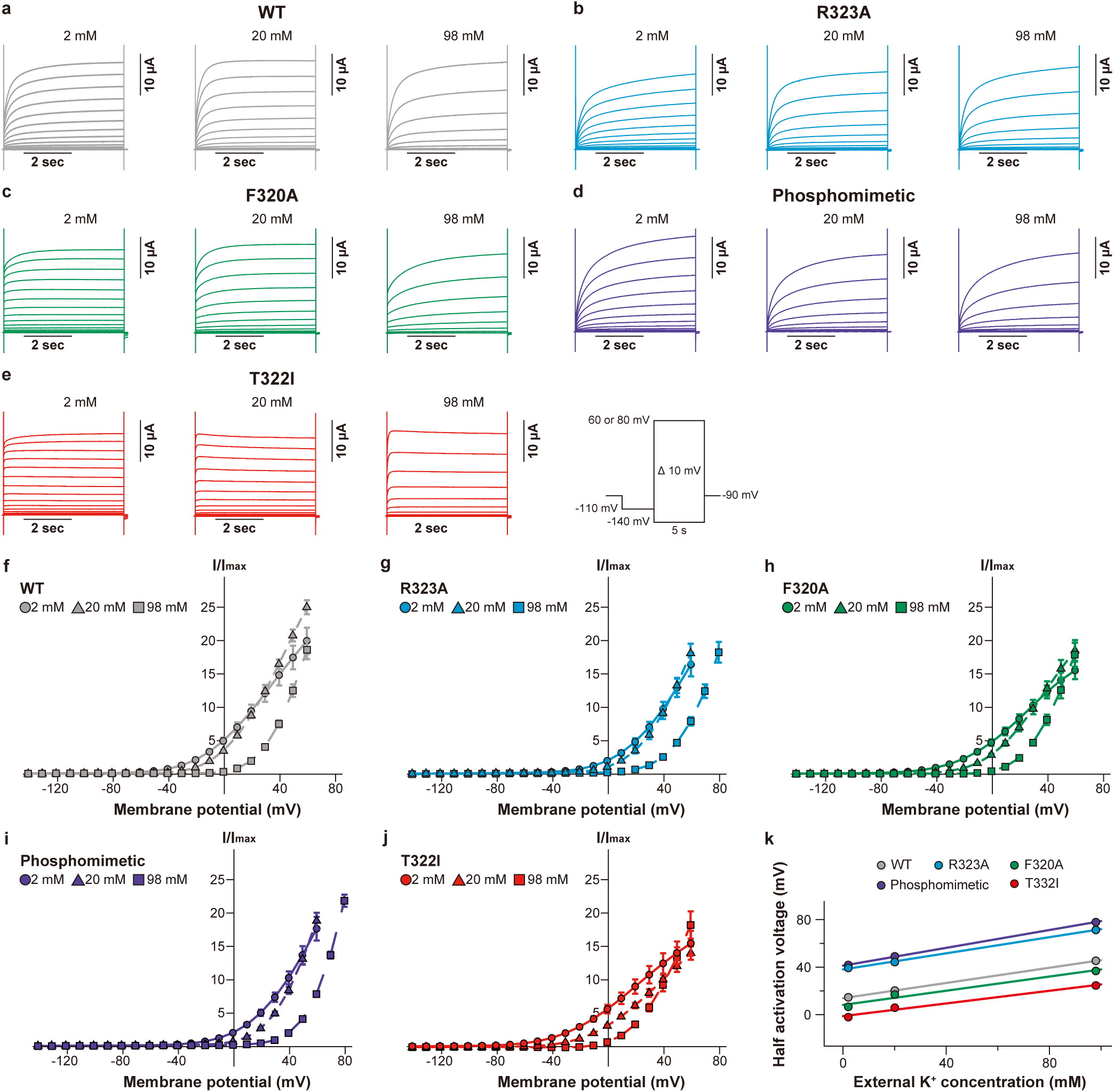
Electrophysiological experiments on TOK1. **a–j.** Representative current traces (**a–e**) and current–voltage (I–V) relationships (**f–j**) of TOK1^WT^ and mutants under external K^+^ concentrations of 2 mM (filled circles), 20 mM (filled triangles), and 98 mM (filled squares). Symbols and error bars represent mean ± s.e.m. (n = 8) (**f–j**). **k.** The relationships between voltage dependence and external K^+^ concentration.

**Supplementary Fig. 6.**
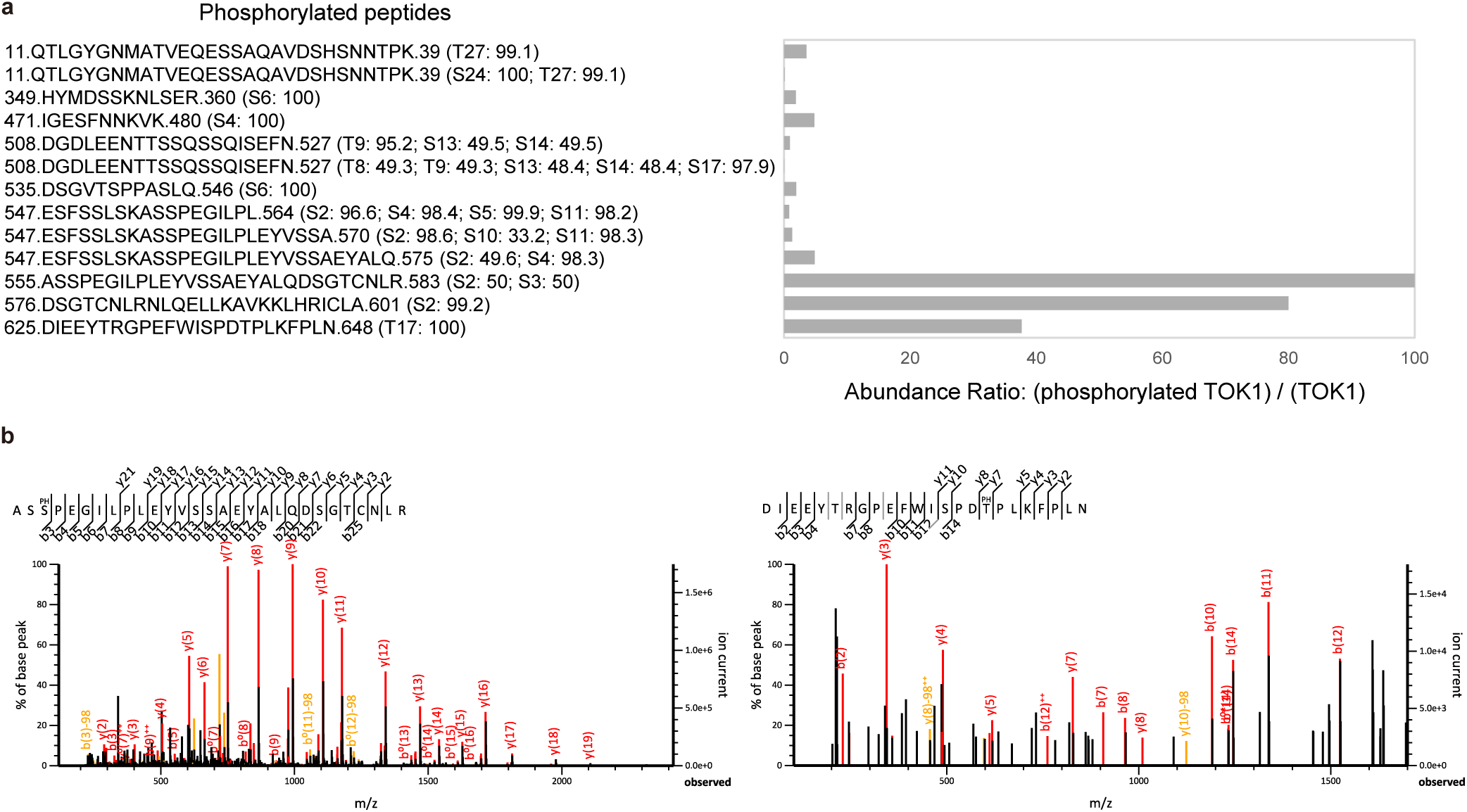
Quantitative and spectral characterization of TOK1-phosphorylated peptides. **a.** Abundance ratios (phosphorylated TOK1 / TOK1) of the identified phosphopeptides. The y-axis indicates peptide sequences, with the corresponding ptmRS Best Site Probabilities shown in parentheses. **b.** Representative MS/MS spectra of phosphopeptides identified by Mascot. The left spectrum was identified with an Ions Score of 113 (Expect = 2.4 × 10^−11^), and the right spectrum with an Ions Score of 26 (Expect = 0.0041).

**Supplementary Fig. 7.**
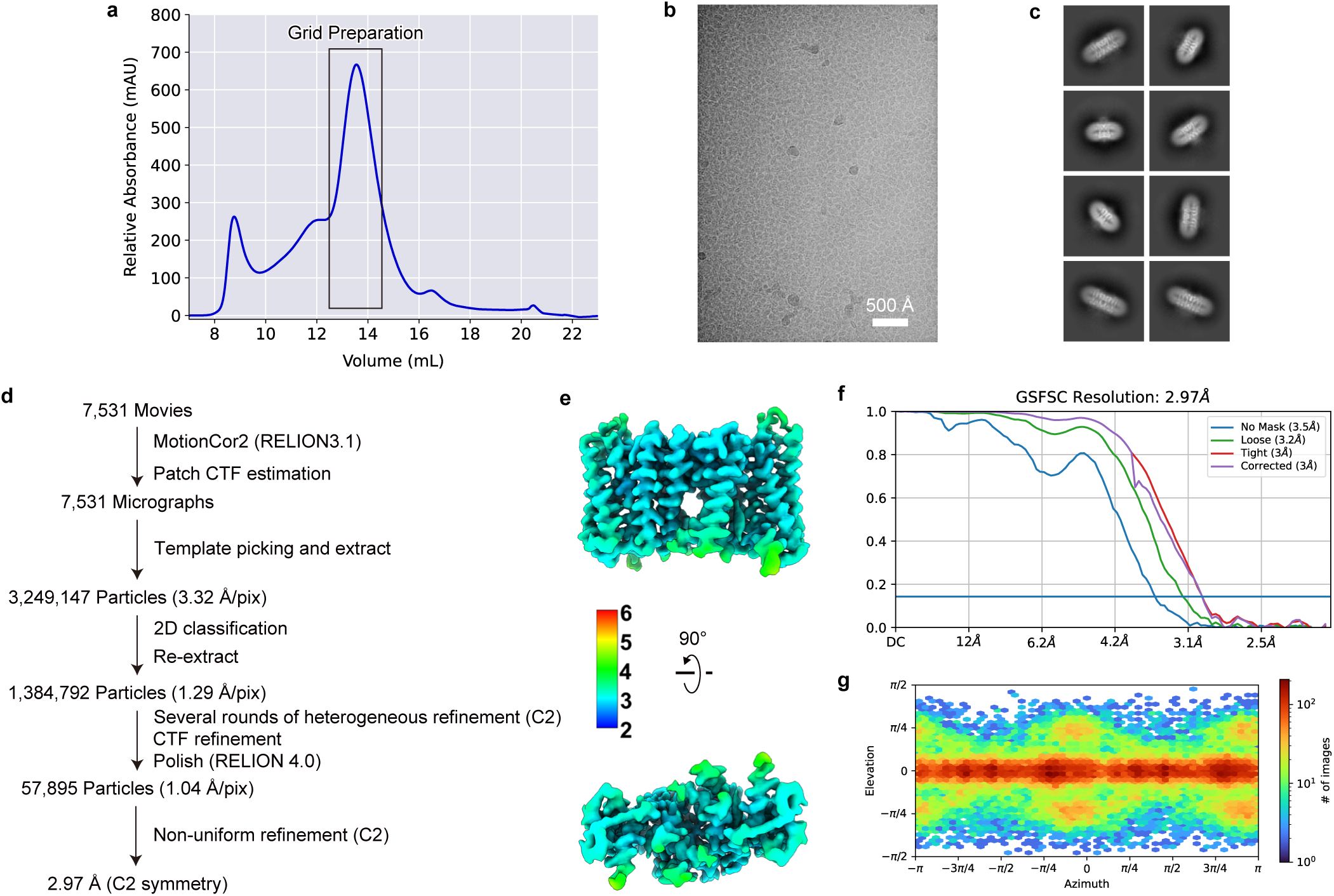
Cryo-EM analysis and map quality of TOK1^phosphomimetic^. **a.** Size-exclusion chromatography profiles of TOK1^phosphomimetic^. **b, c.** Representative cryo-EM micrograph (**b**) and 2D averages (**c**) of TOK1^phosphomimetic^. **d–g.** Cryo-EM workflow (**d**), local-resolution cryo-EM density map (**e**), Fourier shell correlation (FSC) curve (**f**), and angular distribution plots (**g**) of TOK1^phosphomimetic^.

**Supplementary Fig. 8.**
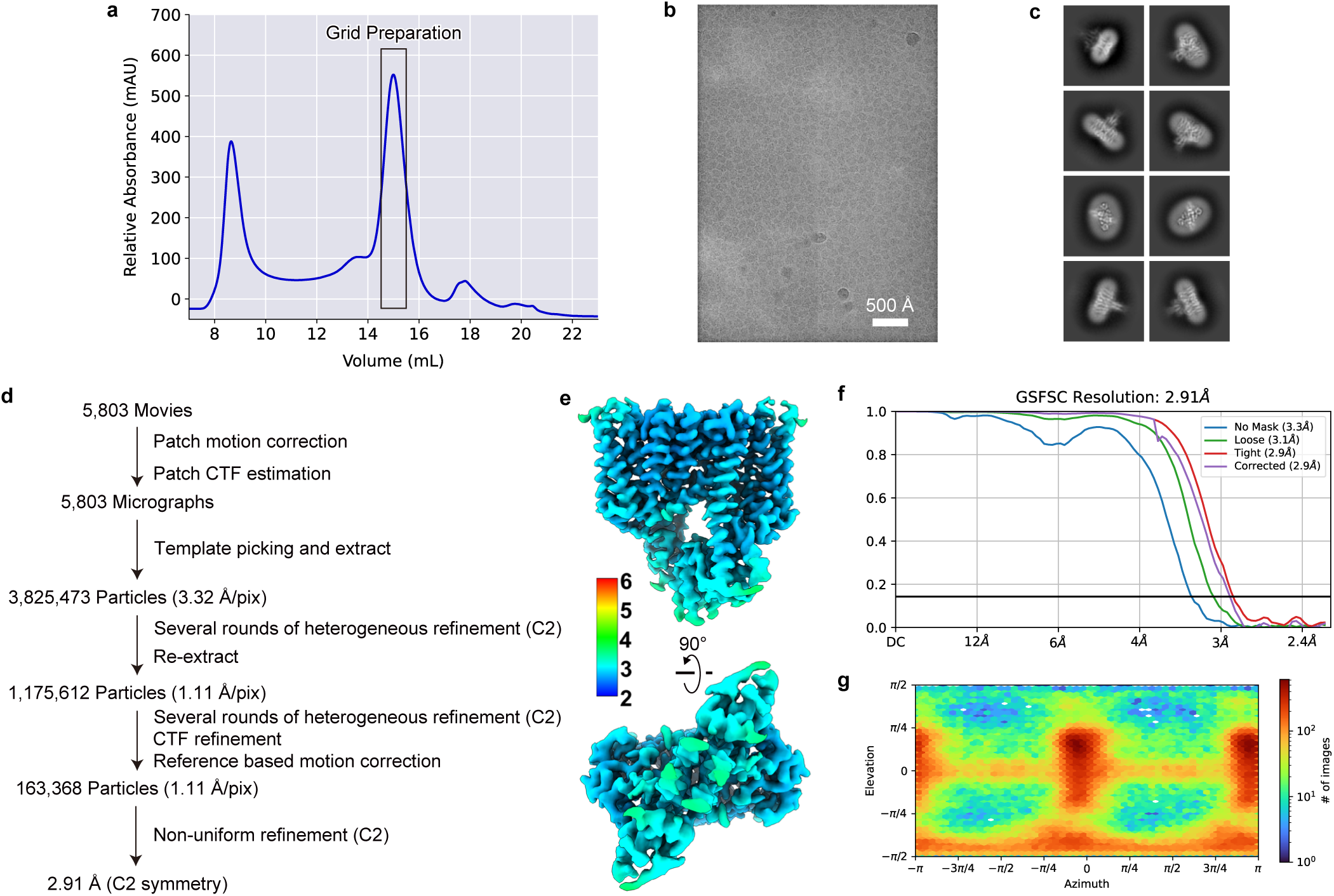
Cryo-EM analysis and map quality of TOK1^T322I^. **a.** Size-exclusion chromatography profiles of TOK1^T322I^. **b, c.** Representative cryo-EM micrograph (**b**) and 2D averages (**c**) of TOK1^T322I^. **d–g.** Cryo-EM workflow (**d**), local-resolution cryo-EM density map (**e**), Fourier shell correlation (FSC) curve (**f**), and angular distribution plots (**g**) of TOK1^T322I^.

**Supplementary Fig. 9.**
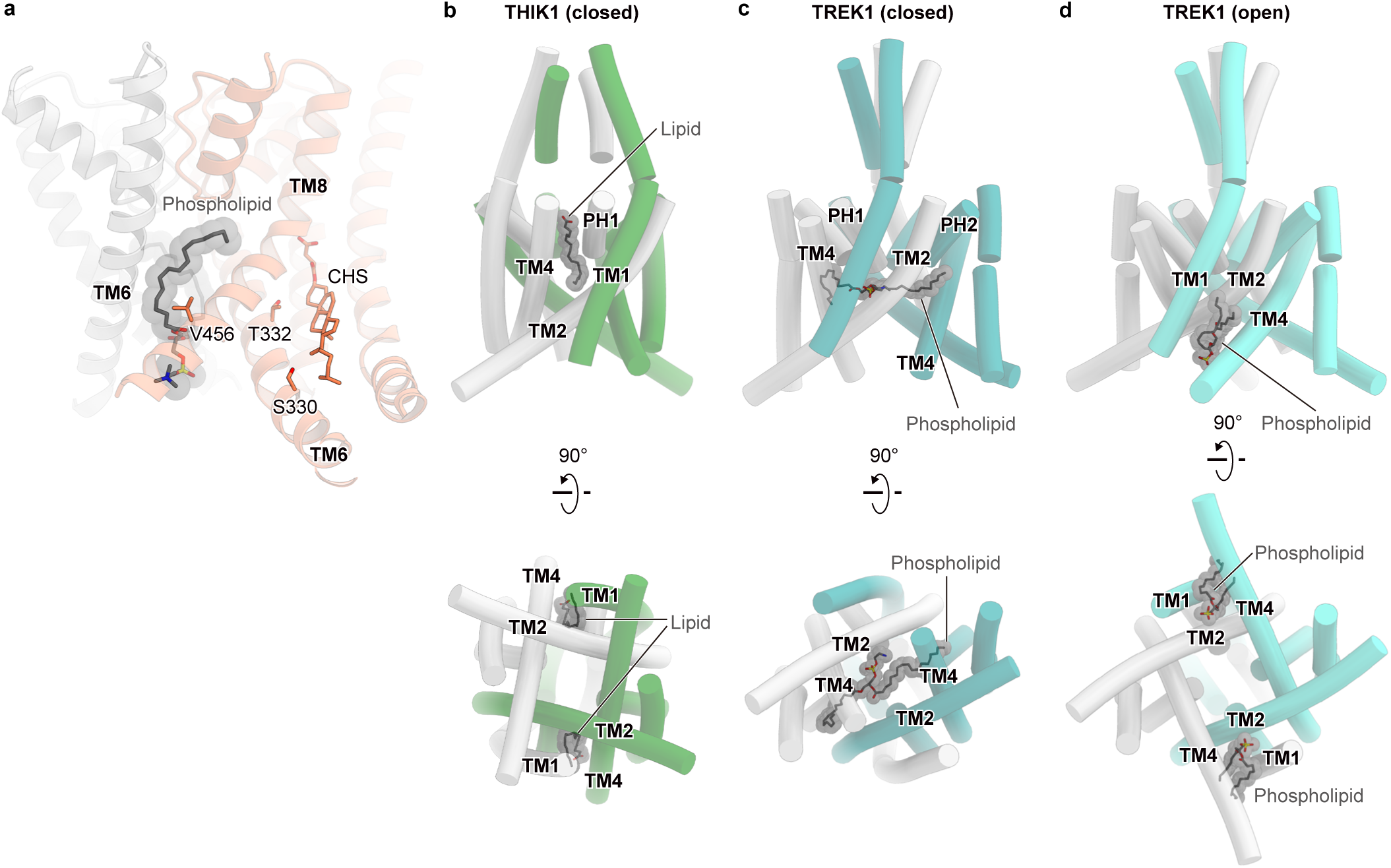
Diverse lipid-binding modes in the K2P channel family. **a.** The S330F mutation likely activates TOK1 by pushing TM8 via CHS, in a manner similar to that of TOK1^T322I^. The V456I mutation may fill the cavity that accommodates the acyl chain of the phospholipid, resulting in lipid dissociation and subsequent channel activation. **b–d.** Lipid-binding modes of THIK1 (closed state, PDB 9FT7, **b**), TREK1 (closed state, PDB 8DE9, **c**), and TREK1 (open state, PDB 8DE8, **d**).

## Supplementary Tables

**Supplementary Table 1.**
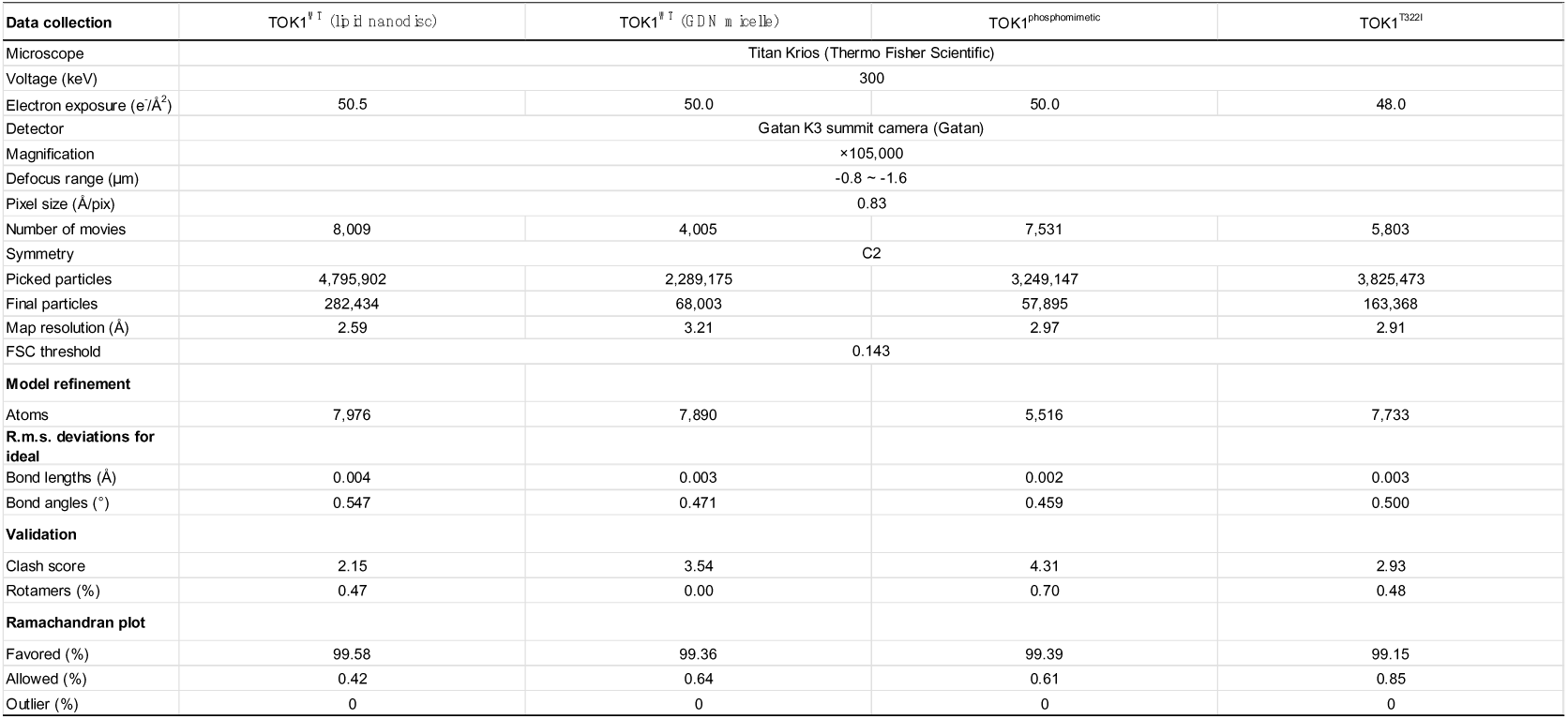
Cryo-EM structural analysis of TOK1.

**Supplementary Table 2.** Electrophysiological recording of TOK1.

| Constructs | External K<sup>+</sup> conc (mM) | *I*<sub>@60 mV</sub> (μA) | *V*<sub>1/2</sub> (mV) | *z* | *n* |
| --- | --- | --- | --- | --- | --- |
| ScTOK1 WT | 2 | 19.9 ± 2.0 | 14.7 ± 1.1 | 1.2 ± 0.0 | 8 |
|  | 20 | 24.9 ± 1.1 | 20.2 ± 1.0 | 1.2 ± 0.1 | 8 |
|  | 98 | 18.6 ± 1.4 | 45.3 ± 0.7 | 1.5 ± 0.0 | 8 |
| ScTOK1 R323A | 2 | 16.3 ± 1.8 | 39.2 ± 1.0 | 1.3 ± 0.0 | 8 |
|  | 20 | 18.0 ± 1.5 | 44.3 ± 1.4 | 1.4 ± 0.0 | 8 |
|  | 98 | 18.2 ± 1.6 (@80 mV) | 71.3 ± 1.0 | 1.4 ± 0.1 | 8 |
| ScTOK1 F320A | 2 | 15.4 ± 1.3 | 5.4 ± 1.0 | 1.2 ± 0.0 | 8 |
|  | 20 | 18.4 ± 1.6 | 16.9 ± 0.5 | 1.2 ± 0.0 | 8 |
|  | 98 | 17.8 ± 1.8 | 36.9 ± 0.8 | 1.5 ± 0.0 | 8 |
| ScTOK1 phosphomimetic | 2 | 17.6 ± 1.8 | 41.8 ± 1.5 | 1.2 ± 0.0 | 8 |
|  | 20 | 18.8 ± 1.2 | 49.0 ± 0.7 | 1.6 ± 0.0 | 8 |
|  | 98 | 21.8 ± 0.9 (@80 mV) | 77.8 ± 1.3 | 1.6 ± 0.0 | 8 |
| ScTOK1 T322I | 2 | 15.4 ± 1.9 | -2.1 ± 1.2 | 1.1 ± 0.1 | 8 |
|  | 20 | 13.9 ± 1.0 | 5.9 ± 1.0 | 1.0 ± 0.0 | 8 |
|  | 98 | 18.1 ± 2.1 | 24.6 ± 1.6 | 1.5 ± 0.0 | 8 |

## Methods

### Constructs

The *Saccharomyces cerevisiae* TOK1 gene was subcloned into the pEG BacMam vector^27^, with an N-terminal twin-Strep tag and C-terminal ALFA^28^ and His_8_ tags. The *Saccharomyces cerevisiae* Hog1 gene was subcloned into the pFastBac vector with an N-terminal His_8_ tag, as previously descrived^29^. The constitutively active version of *Saccharomyces cerevisiae* Pbs2 (S514E, T518E)^30^ was subcloned into the pFastBac vector with an N-terminal His_8_ tag.

### Expression and purification of *Saccharomyces cerevisiae* TOK1

Bacmid preparation and virus production were performed according to the BacMam system (Thermo Fisher Scientific), as previously described^31^.

For the structural analysis of TOK1^WT^ in lipid nanodisc, HEK293S GnTI^−^ (N-acetylglucosaminyl-transferase I-negative) cells (American Type Culture Collection, catalog no. CRL-3022) were cultured in Freestyle 293 medium (Invitrogen) supplemented with 2% FBS (Sigma-Aldrich) in the presence of 8% CO_2_, and infected by P2 baculovirus at a density of approximately 4 × 10^6^ cells ml^−1^. After a 12 hours incubation at 37 °C, the culture was supplemented with 10 mM sodium butyrate and incubated at 30 °C for 65 hours. The cultured cells were collected by centrifugation (5,000*g*, 12 min, 4°C) and disrupted by sonication in lysis buffer (20 mM Tris-HCl pH 8.0, 150 mM KCl, 10% glycerol, 5.2 μg/ml aprotinin, 2.0 μg/ml leupeptin, 1.4 μg/ml pepstatin A, and 100 μM PMSF). The membrane fraction was collected by ultracentrifugation (186,000*g*, 1 hour, 4 °C), and solubilized for 1 hour at 4 °C in solubilization buffer (20 mM Tris-HCl pH 8.0, 150 mM KCl, 10% glycerol, 5.2 μg/ml aprotinin, 2.0 μg/ml leupeptin, 1.4 μg/ml pepstatin A, and 100 μM PMSF, 1% n-dodecyl-beta-D-maltopyranoside (DDM)(Calbiochem), 0.2% cholesteryl hemisuccinate (CHS)(Sigma-Aldrich)). Insoluble materials were removed by ultracentrifugation (142,400*g*, 30 min, 4 °C), and incubated with Strep-Tactin Sepharose resin (IBA) for 1 min. Strep-Tactin Sepharose resin was washed with 20 column volumes of wash buffer (20 mM Tris-HCl pH 8.0, 500 mM KCl, 10% glycerol, 0.03% DDM, 0.006% CHS, 1 mM EDTA), and then eluted with the elution buffer (20 mM Tris-HCl pH 8.0, 150 mM NaCl, 150 mM KCl, 1 mM EDTA, 2.5 mM desthiobiotin, 10% glycerol, 0.03% DDM, 0.006% CHS). The protein was reconstituted in nanodiscs. The TOK1^WT^, MSP2N2, and POPC were mixed at a molar ratio of 1:4:200, respectively, and rotated at 4 °C for 1 hour. Detergents were removed by adding Bio-Beads SM2 (Bio-Rad) to 75 mg ml−1, followed by gentle agitation. Fresh Bio-Beads were added after overnight incubation. The mixture was purified by size-exclusion chromatography on a Superose 6 Increase Column equilibrated in S.E.C. buffer (20 mM Tris-HCl pH 8.0 and 150 mM KCl). The peak fractions were collected and concentrated to an absorbance (A280) of 5.

For the structural analysis of TOK1^WT^ in GDN, the sample was prepared using a protocol similar to that used for TOK1^WT^ in lipid nanodisc. Membrane fraction solubilization was carried out with 1% lauryl maltose neopentyl glycol (LMNG)(Anatrace) and 0.1% CHS, instead of 1% DDM and 0.2% CHS. The Strep-Tactin Sepharose resin was eluted with an elution buffer containing 20 mM Tris-HCl (pH 8.0), 150 mM NaCl, 150 mM KCl, 1 mM EDTA, 2.5 mM desthiobiotin, 10% glycerol, and 0.1% glyco-diosgenin (GDN)(Anatrace). The eluate was purified by size-exclusion chromatography on a Superose 6 Increase Column equilibrated in S.E.C. buffer (20 mM Tris-HCl pH 8.0, 150 mM KCl, 0.03% GDN). The peak fractions were collected and concentrated to approximately 8.8 mg ml^−1^.

For the structural analysis of TOK1^phosphomimetic^, the sample was prepared using a protocol similar to that used for TOK1^WT^ in lipid nanodisc. The eluate was exchanged into GDN detergent and further purified by size-exclusion chromatography on a Superose 6 Increase column equilibrated with SEC buffer (20 mM Tris-HCl pH 8.0, 150 mM KCl, 0.03% GDN). The peak fractions were collected and concentrated to approximately 14.0 mg ml^−1^.

For the structural analysis of TOK1^T322I^, the sample was prepared using a protocol similar to that used for TOK1^WT^ in lipid nanodisc. Membrane fraction solubilization was carried out with 1% LMNG) and 0.1% CHS, instead of 1% DDM and 0.2% CHS. The Strep-Tactin Sepharose resin was eluted with an elution buffer containing 20 mM Tris-HCl (pH 8.0), 150 mM NaCl, 150 mM KCl, 1 mM EDTA, 2.5 mM desthiobiotin, 10% glycerol, and 0.1% GDN. The eluate was purified by size-exclusion chromatography on a Superose 6 Increase Column equilibrated in S.E.C. buffer (20 mM Tris-HCl pH 8.0, 150 mM KCl, 0.03% GDN). The peak fractions were collected and concentrated to approximately 7.2 mg ml^−1^.

### Grid preparation and cryo-EM data collection

The purified TOK1 was applied onto a freshly glow-discharged Quantifoil holey carbon grid (R1.2/1.3, Au, 300 mesh) and blotted at 4 °C in 100% humidity for 4.0 seconds. The grid was then plunge-frozen in liquid ethane, using a Vitrobot Mark IV (FEI). Data collections were performed on a Titan Krios G3 and G4i microscope (Thermo Fisher Scientific) running at 300 kV and equipped with a BioQuantum K3 imaging filter and a K3 direct electron detector (Gatan) in electron counting mode. All movies were acquired at a nominal magnification of 105,000× with a calibrated pixel size of 0.83 ÅLpixel^-1^ using EPU software (Thermo Fisher Scientific) with a defocus range of −0.8 to −1.6 μm. Details are provided in **Supplementary Figs. 2, 3, 7, and 8** and **Supplementary Table 1**.

### Cryo-EM image processing

All acquired movies were dose-fractionated and subjected to beam-induced motion correction using RELION 3.1^32^ or the Patch Motion Correction algorithm implemented in cryoSPARC^33^. The contrast transfer function (CTF) parameters were estimated using the Patch CTF Estimation algorithm. Particles were picked using the Template Picker or Topaz^34^ and subsequently extracted. Then, the particles were subjected to several rounds of 2D classification and heterogeneous refinement to remove ice contamination and poorly aligned particles. The curated particles were further subjected to CTF refinement and Bayesian polishing^35^ or reference-based motion correction. Finally, the particles were refined with C2 symmetry imposed using non-uniform refinement, and the final resolution was estimated according to the Fourier shell correlation (FSC) = 0.143 criterion. Finally, the maps were sharpened and filtered according to the local resolution using the Local Filter tool included in the cryoSPARC package. Details are provided in **Supplementary Figs. 2, 3, 7, and 8** and **Supplementary Table 1**.

### Model building and refinement

The quality of the density map was sufficient for atomic model building. The initial model was obtained from AlphaFold2^36^. The models were first fitted into the density map using the jiggle-fit function in COOT (v0.9.8)^37^, followed by manual adjustment in COOT. The final models were refined using phenix.real_space_refine (v1.20)^38^. The statistics of the 3D reconstruction and model refinement are summarized in **Supplementary Table 1**. All molecular graphics were prepared using CueMol (v2.3; http://www.cuemol.org) and UCSF ChimeraX (v1.10)^39^.

### Molecular dynamics simulation

MD simulations were performed with GROMACS 2020.3^40^. The CHARMM36m force field was used for molecular topology and parameter assignments^41^. For the additional ligands, topology and parameter files were generated using the CGenFF server^42^. The cryo-EM structure of the TOK1^WT^ in lipid nanodisc was embedded into a 1-palmitoyl-2-oleoyl-sn-glycero-3-phosphocholine (POPC) membrane bilayer generated using CHARMM-GUI^43^. The initial system dimensions were 152 × 152 × 130 Å and comprised 562 POPC lipids, 65,792 TIP3P water molecules, 292 chloride ions and 270 potassium ions. The simulation systems were energy-minimized until the maximum force was reduced below 1,000 kJ mol^−1^ nm^−2^ with fixed positions of the nonhydrogen atoms. After minimization, equilibration were performed under NVT and NPT ensembles for 1 and 10 ns, respectively, with 1,000 kJ mol^−1^ nm^−2^ restraints for heavy atoms of the protein. The production runs of the equilibrium simulations were performed three times for each 250 ns without any restraints. The detailed conditions for the production runs are as follows: a 2 fs timestep in the NPT ensemble with semi-isotropic coupling at 310 K and 1 bar, maintained by the Nose-Hoover thermostat and Parrinello–Rahman barostat. The long-range electrostatic interactions were calculated by the particle mesh Ewald method. The simulation results were analyzed using PyMOL (version 2.5.0; https://pymol.org/2/), GetContacts (https://getcontacts.github.io/), and MDTraj (version 1.9.7)^44^.

### Protein expression in *Xenopus laevis* oocytes

The *Saccharomyces cerevisiae TOK1* (WT and mutants) cDNAs were inserted into the pGEMHE expression vector^45^. The cRNAs were transcribed using the mMESSAGE mMACHINE^TM^ T7 Transcription Kit (ThermoFisher Scientific, AM1344). Oocytes were surgically extracted from female *Xenopus laevis* and anesthetized in water containing 0.15% tricaine (Sigma-Aldrich, E10521) for 15-30 minutes. They were treated with collagenase (Sigma-Aldrich, C0130) at room temperature for 5-6 hours to remove the follicular cell layer. Defolliculated oocytes of a similar size at stage V or VI were selected for cRNA injection. The 50 nL of cRNA solution (2-20 ng) was injected into each oocyte using Nanoject II (Drummond Scientific Company). cRNA-injected oocytes were incubated for 1-4 days at 18 °C in MBSH buffer (88 mM NaCl, 1 mM KCl, 2.4 mM NaHCO_3_, 10 mM HEPES, 0.3 mM Ca(NO_3_)_2_, 0.41 mM CaCl_2_, and 0.82 mM MgSO_4_, pH 7.6) supplemented with 0.1% penicillin-streptomycin solution (Sigma-Aldrich, P4333). All experiments were approved by the Animal Care Committee of Jichi Medical University under protocol no. 21030-03 and were performed following the guidelines.

### Two-electrode voltage clamp

Microelectrodes were drawn from borosilicate glass capillaries (Sutter Instruments, B150-117-10) using a P-1000 micropipette puller (Sutter Instrument) to achieve a resistance of 0.2-1.0 MΩ and filled with 3 M KCl. The bath solutions used were: 2 mM KCl solution (ND96 solution) (96 mM NaCl, 2 mM KCl, 1.8 mM CaCl_2_, 1 mM MgCl_2_, and 5 mM HEPES, pH 7.5); 20 mM KCl solution (78 mM NaCl, 20 mM KCl, 1.8 mM CaCl_2_, 1 mM MgCl_2_, and 5 mM HEPES, pH 7.5); and 98 mM KCl solution (KD98 solution) (98 mM KCl, 1.8 mM CaCl_2_, 1 mM MgCl_2_, and 5 mM HEPES, pH 7.5). Voltage-clamp protocols were generated and data were acquired using a Digidata 1550 interface (Molecular Devices) controlled by pCLAMP 10.7 software (Molecular Devices). Data were sampled at 10 kHz and filtered at 1 kHz. Details are provided in **Supplementary Table 2**.

### Data analysis of electrophysiological experiments

The holding potential was set at -90 mV. After 0.8 seconds of hyperpolarization at -110 mV, currents were elicited by 5-second test pulses to membrane potentials from -140 to 60 mV or -140 to 80 mV with 10-mV increments. Conductance values were calculated from peak current amplitudes by normalizing to the maximum current amplitude obtained in the experiment, assuming a linear open channel current-voltage relationship (normalized chord conductance). The reversal potential for each K^+^ condition was calculated, assuming an intracellular K^+^ concentration of *Xenopus* oocytes was 108 mM, which is the average value of Table 2 in the previous report^46^. Normalized peak conductance was plotted versus voltage and fitted with single Boltzmann functions to estimate the half-activation voltage (*V_1/2_*) and the effective charge (*z*). Oocytes with a holding current larger than -0.4 μA at -90 mV were excluded from the analysis. Oocytes with a maximum current amplitude at the most positive voltage in the recording (60 mV or 80 mV) between 10 µA and 30 µA were used for the analysis. Details are provided in **Supplementary Table 2**.

### Statistical analysis of electrophysiological experiments

The electrophysiological data were expressed as mean ± s.e.m (*n* = 8). Details are provided in **Supplementary Table 2**.

### Mass spectrometry

For the expression and purification of Pbs2 and Hog1, bacmid preparation and virus production were performed according to the Bac-to-Bac system (Gibco, Invitrogen), as previously described^47^. Hog1 and Pbs2 were expressed and purified separately following the procedures described below. *Spodoptera frugiperda* (Sf9; Gibco) cells at a density of 3 × 10^6^ cells ml^-1^ were infected with P2 baculoviruses. The infected Sf9 cells were incubated in Sf900II medium at 27 °C for 48–60 hours. The Sf9 cells were collected by centrifugation (5000*g,* 12 min, 4°C), and disrupted by sonication in lysis buffer (20 mM Tris-HCl pH 8.0, 150 mM NaCl, 10% glycerol). For Hog1, the buffer was further supplemented with 5.2 μg/ml aprotinin, 2.0 μg/ml leupeptin, 1.4 μg/ml pepstatin A, and 100 μM PMSF. Insoluble materials were removed by ultracentrifugation (142,400*g* or 186,000*g*, 1 hour, 4 °C), and incubated with Ni-NTA resin (Qiagen) for 1–6 hours. Ni-NTA resin was washed with 15–25 column volumes of wash buffer (20 mM Tris-HCl pH 8.0, 500 mM NaCl, 10% Glycerol, 30 mM imidazole), and then eluted with the elution buffer (20 mM Tris-HCl pH 8.0, 150 mM NaCl, 10% Glycerol, 300 mM imidazole). The eluate was further purified by size-exclusion chromatography on a Superdex 200 10/300 Increase column equilibrated in S.E.C. buffer (20 mM Tris-HCl pH 8.0, 150 mM NaCl). The peak fractions were collected and concentrated to approximately 8 mg ml^−1^. The protein was frozen in liquid nitrogen after supplemented 10% of glycerol.

For the purification of TOK1, the sample was prepared using a protocol similar to that used for the structural analysis of TOK1^WT^ in lipid nanodisc. The eluate was further purified by size-exclusion chromatography on a Superose 6 Increase column equilibrated with SEC buffer (20 mM Tris-HCl pH 8.0, 150 mM KCl, 0.03% DDM, 0.006% CHS). The peak fractions were collected and concentrated to approximately 1.3 mg ml^−1^. The protein was frozen in liquid nitrogen after supplemented 10% of glycerol.

A total of 375 μg of purified Pbs2 was mixed with 750 μg of purified Hog1 in kinase buffer (50 mM Tris-HCl pH 8.0, 10 mM MgCl_2_, 50 mM ATP, 0.15% DDM, and 0.03% CHS) and incubated at 37 °C for 1 hour. After 1 hour of incubation, 2.5 mg of purified TOK1 were supplemented, and incubated at 37 °C for 1 hour. The mixture were purified by size-exclusion chromatography on a Superose 6 Increase column equilibrated in S.E.C. buffer (20 mM Tris-HCl pH 8.0, 150 mM KCl, 0.03% DDM, 0.006% CHS). The peak fractions were collected and concentrated to approximately 2.1 mg ml^−1^. The protein was frozen in liquid nitrogen after supplemented 10% of glycerol.

TOK1 and phosphorylated TOK1 were digested with either trypsin (TPCK-treated; Worthington Biochemical Corporation, Lakewood, NJ, USA) or endoproteinase Asp-N (Roche Diagnostics, Mannheim, Germany) at 37 °C for 12 h. The resulting peptides were analyzed by nanoLC–MS/MS using a Q Exactive HF-X Hybrid Quadrupole-Orbitrap mass spectrometer (Thermo Fisher Scientific, Bremen, Germany) coupled to an Easy-nLC 1200 system (Thermo Fisher Scientific). Solvent A consisted of 0.1% formic acid in water, and solvent B consisted of 0.1% formic acid in 80% acetonitrile. Peptides were separated on a nano-ESI analytical column (NTCC-360; 0.075 mm i.d. × 150 mm, 3 µm particle size; Nikkyo Technos Co., Ltd., Tokyo, Japan) at a flow rate of 300 nL/min using a linear gradient from 0% to 80% solvent B over 20 min. The eluted peptides were ionized using a nano-electrospray ionization source and analyzed in positive ion mode with data-dependent acquisition (DDA) using the Top20 method. The acquired MS/MS spectra were processed using Mascot 3.1 (Matrix Science, London, UK) and Proteome Discoverer 3.1 (Thermo Fisher Scientific). Database searches were performed against the SwissProt database (taxonomy: Saccharomyces cerevisiae) with the following parameters: enzyme specificity, trypsin or Asp-N; maximum missed cleavages, 4; variable modifications, acetylation (protein N-terminus), pyroglutamate formation from N-terminal Gln, oxidation (Met), and phosphorylation (Ser/Thr); peptide mass tolerance, ±15 ppm; fragment mass tolerance, ±30 mmu; instrument type, ESI-TRAP. Phosphorylation site localization probabilities were determined using the ptmRS node implemented in Proteome Discoverer. Quantification was performed using the label-free quantification (LFQ) workflow implemented in Proteome Discoverer 3.1.

